# Connecting Brain Responsivity and Real-World Risk taking: Strengths and Limitations of Current Methodological Approaches

**DOI:** 10.1101/142331

**Authors:** Lauren Sherman, Laurence Steinberg, Jason Chein

**Affiliations:** Department of Psychology Temple University

## Abstract

In line with the goal of limiting health risk behaviors in adolescence, a growing literature investigates whether individual differences in functional brain responses can be related to vulnerability to engage in risky decision-making. We review this body of work, investigate when and in what way findings converge, and provide best practice recommendations. We identified 23 studies that examined individual differences in brain responsivity and adolescent risk taking. Findings varied widely in terms of the neural regions identified as relating to risky behavior. This heterogeneity is likely due to the abundance of approaches used to assess risk taking, and to the disparity of fMRI tasks. Indeed, brain-behavior correlations were typically found in regions showing a main effect of task. However, results from a test of publication bias suggested that region of interest approaches lacked evidential value. The findings suggest that neural factors differentiating riskier teens are not localized to a single region. Therefore, approaches that utilize data from the entire brain, particularly in predictive analyses, may yield more reliable and applicable results. We discuss several decision points that researchers should consider when designing a study, and emphasize the importance of precise research questions that move beyond a general desire to address adolescent risk taking.

---

A stated goal of much research on the neural bases of adolescent decision-making has been to address the significant public health problems thought to result from adolescent risk taking, including alcohol and drug abuse, automobile accidents, and unprotected sex (e.g., Bjork and Pardini, 2015; Chein, Albert, O’Brien, Uckert, and Steinberg; Galvan et al., 2006; Pfiefer et al., 2011). Two prominent theoretical papers about adolescent decision-making open with a description of these reckless behaviors, and propose that a neurobiological framework may account for the particular susceptibility of adolescents to risky behavior (Casey, Getz, and Galvan, 2008; Steinberg, 2008). In 2008, Steinberg proposed the “Dual Systems Model,” which postulates that the differing trajectories of two systems in the brain, a “socio-emotional system” and a “cognitive control system,” make mid-adolescence a period of heightened vulnerability. The “Imbalance Model,” discussed in Casey, Getz, and Galvan’s 2008 paper, also proposes that the developmental trajectories of two brain systems, implicated in reward response and top-down control, respectively, confer heightened vulnerability to reckless behavior in adolescence.

Work over the past nine years has led to substantiation as well as refinement of these models (e.g. Casey, Galvan, and Somerville, 2016; Shulman et al., 2016), including a braod acknowledgment that risk taking in adolescence can be, and often is, adaptive (e.g. Dahl, 2016). Nonetheless, the concerns cited in 2008 represent ongoing public health issues. Health risk behaviors are still leading causes of mortality, permanent injury, and other problems among adolescents (Centers for Disease Control and Prevention, 2015). Furthermore, some health risk behaviors, such as texting and driving and the use of electronic cigarettes, have only recently been identified as especially worrisome among adolescents (Centers for Disease Control and Prevention, 2015). Thus, if neurobiological models of decision-making can provide additional insight into the causes underlying adolescent health risk behaviors, above and beyond behavior alone, an important next step would be leveraging this knowledge of mechanisms to develop targeted preventative measures.

One potential avenue for the improvement of preventative measures is through the identification of individual differences in vulnerability to risk taking among adolescents. If heightened adolescent risk taking can be explained by processes taking place within the adolescent brain in a way that is different from adults and children, is it also the case that adolescents with a *higher* predilection for risk will show exaggerated brain differences? If individual differences in adolescent risk taking *can* be predicted by brain responses, then the brain systems implicated in reward and cognitive control would be sensible targets of investigation. However, the leap from age differences to individual differences is not a foregone conclusion. It may be, for example, that the variability of brain responses *within* the adolescent period is not sufficient to predict behavioral outcomes among people this age, or that brain regions predictive of outcomes within a single age band are distinct from those that are correlated with age.

As noted by Pfeifer and Allen (2013), early neuroimaging work on adolescent decisionmaking rarely considered individual differences in brain responses as they relate to behavior. More recently, Bjork and Pardini (2015) argued that the maturational imbalance proposed in neural models of adolescent risk taking might actually be more characteristic of individuals with histories of excessive behavioral disinhibition, and suggested that, “Future research on brain mechanisms of significant adolescent risk taking would therefore benefit from a greater emphasis on individual differences.”

Since the publications of these papers, developmental cognitive neuroscientists have increasingly asked how individual differences in neural responses are related to risk-taking behavior in the laboratory and beyond, and have been successful in identifying significant correlations between patterns of neural activity and aspects of risk-related behavior. The goal of the present review is to provide a systematic evaluation of this body of work and to address two overarching questions. First, do individual difference findings converge to suggest that neural markers of individual adolescent vulnerability for risk mirror factors that have been identified in studies of broader developmental brain changes? And second, what do these findings teach us about best practices moving forward?

## Approach

### Definitions and inclusion criteria

In line with our stated goal of identifying the neural markers of vulnerability in adolescence, we restricted our review to studies that met the following criteria: they examined *individual differences* in risk-taking behaviors *in adolescent samples* in relation to responses *during task-based fMRI*. We defined individual differences broadly, as variability in any continuous measures of risky behavior. “Risky behavior” was defined as either lab-based risky decision-making (e.g. riskier behavior in a gambling game), reports of health risk behaviors such as alcohol and drug use, or high scores on personality measures that have been implicated in adolescent risk taking, such as sensation-seeking (e.g. Steinberg, 2008). Our motivation for casting this wide net was twofold: first, we hoped to assemble a critical mass of papers to perform quantitative analyses as well as qualitative description; and second, we wanted our review to reflect the wide range of operational definitions that researchers have used when attempting to understand neural underpinnings of risk taking and risky decision-making in adolescence. It is important to note, however, that the term ‘risk’ is defined differently across different literatures; economists and neuroeconomists typically define risk in terms of the uncertainty of possible outcomes, whereas clinicians and developmental cognitive neuroscientists often define risk as behaviors that confer potential harm (Defoe, et al., 2014; Schonberg, Fox, and Poldrack, 2011).

In this review, we use the term “individual differences” to refer to psychological phenomena that vary continuously (e.g. Dubois and Adolphs, 2016), and focus our review on papers that utilize correlational or regression approaches, rather than group comparisons. This approach is now fairly common in the developmental cognitive neuroscience literature, and reflects the assumption that the phenomenon of risk taking varies within the population of typical adolescents. It should be noted that a complementary and well-established clinical literature mainly has taken an alternative, categorical, approach of comparing structural and functional differences between youth with versus without substance use disorders (for two recent reviews, see Heitzeg, Cope, Martz, and Hardee, 2015 and Squeglia and Cservenka, 2017). In contrast to that work, we wished to establish whether risk taking, conceptualized as a continuously variable phenomenon within the general population rather than an aberrant outcome, is related to differences in brain responses. Further, we hoped to make specific methodological recommendations for correlational and regression approaches. As we we later discuss, however, the use of a correlational or categorical approach is a decision that should be based on the goals of the individual study, hypothesized mechanisms, and one’s stance regarding the value of work in neurotypical youth to ameliorate the public health consequences of risky behavior.

Like “risk taking,” the term “adolescence” has been defined a number of ways, not only across disciplines, but within developmental cognitive neuroscience (Crone and Dahl, 2012). Frequently, it is defined as beginning with pubertal onset (e.g., Graber & Brooks-Gunn, 1998) and ending with independence from parents (e.g., Casey, Duhoux, and Cohen, 2010), though these markers, of course, are not precise moments in time but rather transitions in and of themselves. Many of the neural changes hypothesized to underlie the rise in risky behaviors occur during the second decade of life (Steinberg, 2008; Casey et al., 2008), but these processes too are continuous and not restricted to the teen years (e.g. changes in grey and white matter volume, Mills et al., 2016; changes in reward sensitivity in the ventral striatum; Braams et al., 2015). In the present review, we included any papers for which the mean age, as well as the age of the majority of participants, fell within the second decade of life. As with our definition of “risk taking,” we used an intentionally liberal boundary for age range in order to maximize the size of the reviewed literature.

This review does not explore the relation between neural response and age differences in risk taking as a function of *age* or *developmental stage*; that is, we do not review the literature that compares adolescents to adults and/or children in terms of risky behavior and brain development. For more information on this topic we refer readers to two extensive recent metaanalyses, the first on reward processing in adolescence (with a comparison to adulthood; Merav, Jedd, and Luciana, 2015), and the second, on differences in laboratory risk taking between children, adolescents, and adults (Defoe et al., 2014). While dual systems and imbalance models propose a direct relation between the development of reward circuitry and risk-taking behavior, these two phenomena are confounded with age and other related variables (e.g., cohort, years of experience); here, instead, we look at brain-behavioral relations *within the developmental window of adolescence*. Several studies in the literature separate their samples into adolescents, adults, and/or children; in these cases, we used only the adolescent sample as defined by the authors. These cases are noted in Table 1.

**Table 1.** Studies investigating brain-behavior correlations related to adolescent risk-taking.

| Authors (organized by fMRI task category) | Measure of risk-taking | Task | A priori ROI? | Whole brain / bottom up? | Regions in whole brain correlation | n | Ages | Longitudinal? |
| --- | --- | --- | --- | --- | --- | --- | --- | --- |
| **Monetary Incentive Delay** |  |  |  |  |  |  |  |  |
| Bjork et al., 2011 | Overall Problem Density on the Drug Use Screening Inventory | MID | VS (sig) | Yes | mPFC | 26 | 12 to 17 | No |
| Bjork, et al., 2008 | Brief Sensation-Seeking Scale | MID | NACC (sig) | No |  | 26 | 12 to 16 | No |
| Benningfield, et al., 2014 | Delay discounting behavior during task (Monetary Choice Questionnaire) | MID | No | Yes, within task activation mask | left ventromedial caudate | 19 | 10 to 14 | No |
| Nees et al., 2012 | Alcohol Use Disorders Identification Test (ns), European School Survey Project on Alcohol and Drugs (ns), Fagerstrom Test for Nicotine Dependence (ns); Novelty-seeking scale of the Temperament and Character Inventory-Revised (ns); Substance Use Risk Profile Scale (ns); Risk-taking on the Cambridge Gambling Task (CGT; sig) | MID | Insula, amygdala, PFC, OFC, NAcc, ACC, putamen, nucleus caudatus, thalamus, cerebellar vermis. Only significant bivariate correlations were that putamen and thalamus correlated with risk-taking on CGT. All brain regions were used in SEM analysis | No |  | 324 | 14 | No |
| **Other Reward Tasks (no risky decision-making)** |  |  |  |  |  |  |  |  |
| Braams et al., 2016 | Reported alcohol use: drinks per day (sig) and lifetime alcohol use (ns) | Coin-flip reward task | NACC (sig) | No |  | 169 | 10 to 26 | Yes |
| Galvan et al., 2007 | Cognitive Appraisal of Risky Events (sig), modified Benthin Risk Perception Measure (ns), Connor's Impulsivity Scale (ns) | Two-choice reward task | NACC (sig) | No |  | 7* | 13 to 17 | No |
| **Stoplight/ Risky Driving Task** |  |  |  |  |  |  |  |  |
| Kahn et al., 2016 | Sensation-Seeking Scale (sig), Sensitivity to Punishment and Sensitivity to Reward- Short Form (sig), Composite of Risk-taking (CRT; ns), Barrett Impulsivity Scale (ns); task riskiness (ns) | Risky driving task | No | Extracted parameters from task activation clusters | IFG (sig); VS (ns) | 20 | 14 to 16 | No |
| Vorobyev et al., 2015 | Task performance on the risky driving task. | Risky driving task | No | Yes | mPFC | 27 | 18 to 19 | No |
| **Gambling Task (Wheel of Fortune/ Cake gambling)** |  |  |  |  |  |  |  |  |
| Cservenka, et al., 2016 | Drinks per day (sig); marijuana use (ns); cigarette use (ns); categorical analysis also performed (group comparison of alcohol naive vs. drinkers) | Wheel of Fortune | No | Extracted parameters from task activation cluster | Cerebellum | 34 | 12 to 16 | Yes |
| Eshel et al., 2007 | Risk taking behavior on the fMRI task (Wheel of Fortune) | Wheel of Fortune | bilateral OFC (left sig, right ns), bilateral ACC (sig) | No |  | 16* | 9 to 17 | No |
| Van Leijenhorst et al., 2010 | Percent high-risk gambles in game | Cake Gambling Task | No | Yes | vmPFC, MTG, IFG, STG, Precuneus, PCC, cingulate, OFC, insula, lateral occipital cortex, BG, frontal pole, middle frontal gyrus, occipital pole | 58 | 8 to 26 | No |
| **Gambling Task (Other)** |  |  |  |  |  |  |  |  |
| Barkley-Levenson et al., 2013 | Domain Specific Risk-Taking Scale | Mixed gambles task | No | Yes | mPFC | 18* | 13-17 | No |
| Op de Macks et al., 2016 | Risk Questionnaire, task-related risk-taking | Jackpot task | NACC (tested for task-related risk-taking only) | Yes | medial OFC | 58 | 11 to 13 | No |
| Paulsen et al., 2012 | Risk aversion during task | Risky gambling task | No | Yes | medial frontal gyrus, central cingulate gyrus | 17* | 14 to 16 | No |
| Xiao et al., 2013 | Drinks in past 30 days (ns), age of alcohol onset (ns), Rutgers Alcohol Problem Index (sig), and urgency score on the UPPS Impulsive Behavior Scale (sig); group comparison between binge drinkers and never drinkers was also performed | Iowa Gambling Task | OFC (neg with drinking problems and urgency), insula (pos with drinking problems and urgency) | No |  | 28 | 16 to 18 | No |
| **Balloon Analogue Risk Task** |  |  |  |  |  |  |  |  |
| McCormick and Telzer, 2017 | Behavioral performance on the BART (differential sensitivity to positive and negative trials); modified version of the Adolescent Risk-Taking Scale | BART | No | Yes | For behavioral measure: mPFC, left dlPFC, insula, caudate; for self-report: mPFC, caudate, pre-SMA | 58 | 12 to 18 | No |
| Telzer et al., 2015 | Behavioral performance on the BART; modified version of the Adolescent Risk-Taking Scale | BART | Based on previous bottom-up analysis with non-risk variable: VS (sig) and insula (sig) | No |  | 46 | Mean age 16.3 | No |
| **Executive Function Tasks** |  |  |  |  |  |  |  |  |
| Cascio et al., 2015 | Behavior during driving task | Go/No-Go | BG and rIFG. Significant interaction with peer manipulation. In the presence of a cautious, but not risky peer, greater BG/IFG activation during Go/No-Go associated with safer driving behavior. | Yes | vmPFC, putamen, caudate, mtg, fusiform, hippocampus, paracentral lobe, occipital lobe (association was with cautious peers but not risky peers) | 37 | 16 to 17 | No |
| Chung et al., 2015 | Marijuana symptom count from DSM-IV SUDs, adapted for adolescents | Reward cue anti-saccade task | Amygdala (sig), caudate (ns), NAcc (sig), OFC (ns), putamen (sig), vmPFC (ns), dACC (ns), dlPFC (ns), FEF (ns), PPC (ns), preSMA (ns), SEF (sig), vlPFC (sig) | No |  | 14 | 14 to 18 | Yes |
| Mahmood et al., 2013 | Customary Drinking and Drug Use Record], Fagerstrom Test for Nicotine Dependence | Go/No-Go | No, but only regions from task activation contrast were used in regression model | Yes, for initial task activation map, which was then submitted to regression analysis | Angular gyrus pos correlation with drug use and vmPFC neg correlation with drug and alcohol dependency; high-frequency users only | 80 | 16 to 19 | Yes |
| **Other** |  |  |  |  |  |  |  |  |
| Pfeifer et al., 2011 | Risk-taking Questionnaire; questions derived from the Lerner 4-H study of Positive Youth Development | Affective face processing | VS and vmPFC boundaries defined by task, amygdala defined by anatomy. VS significantly negatively related to risk-taking (others not reported, assumed to be NS) | No |  | 38 | Age 10, Age 13 | Yes |
| Saxbe et al., 2015 | Youth Risk Behavior Survey, Peer Behavior Inventory | Emotion rating task with video clips | amPFC (sig), rACC (ns) | Yes | Precuneus, PCC, mPFC | 22 | 15 to 19 | No |
| Sherman et al., 2017 | Revised Cognitive Appraisal of Risky Events | Social media task with risky stimuli | No | Yes, within task-activation map | occipital cortex, precuneus/PCC | 58 | 13 to 21 | No |

Finally, we restricted our review to studies employing task-based functional MRI. This allowed us to make recommendations for best practices within one methodological domain and to quantitatively aggregate the data. As will become clear, even with this somewhat narrow definition, the methods utilized are highly heterogeneous. Task-based fMRI is only one of several categories of approaches that have been used to examine individual differences in adolescent risk taking; others include investigations of cortical volume (e.g., Medina et al., 2009; DeBellis et al., 2005), white matter integrity (e.g., Bava et al., 2009, 2010; McQueeney et al., 2009), and resting-state functional connectivity (e.g., DeWitt, Aslan, and Filbey, 2014; Peters et al., 2015; van den Bos, et al., 2015). Furthermore, while we will briefly discuss measures of connectivity during task-based fMRI, we focus primarily on activation rather than connectivity. The motivation for primarily discussing activation was that this approach dominates in the literature; however, as we will discuss, connectivity and network approaches may yield additional insights, and will likely deserve greater attention as the number of studies adopting connectivity approaches grows.

### Literature search

We relied on several methods to compile the relevant corpus of studies. Initially, we conducted searches on PubMed using the combination “adolescent,” “individual difference,” “risk,” and “fMRI;” this search yielded 32 entries. We substituted the term “risk” with “alcohol,” (6 additional entries) “marijuana,” (1 additional entry) and “sensation-seeking” (no additional entries). The first author reviewed all titles and abstracts and included papers that met the above criteria. Additional papers were identified by reviewing the Introductions and reference lists of the papers identified during the search, as well as through the authors’ prior knowledge of recent literature, and through consultation with other researchers (e.g. by soliciting papers on the Social and Affective Neuroscience Society listserv and via discussion through social media).

In several cases, it was determined that the same sample was used in more than one publication. In these cases, we included the study with the largest sample size in our main table and quantitative review, and noted (in Supplemetary Table 1) which related works were excluded. One exception is the inclusion of both Pfeifer et al. (2011) and Sherman et al., (2017). In this case, only five individuals (out of 38 and 58 total participants, respectively) participated in both studies, with a 4–5 year gap between the two studies. We elected to include both papers because they utilized different approaches to both the fMRI task and outcome measures, and because Sherman et al. was not included in the quantitative analysis; nonetheless, these studies’ nonindependence should be noted.

### Organization and reporting of findings

Our aim was not only to review the literature but also to provide suggestions for best practices. Initially, we were most interested in determining if and how results might differ as a function of the fMRI task used. Thus, we categorized findings in our table by fMRI task, and provide below an overview of the tasks used to assess these brain-behavior relations, while considering advantages and limitations of each approach. Although we initially hoped to perform quantitative analyses comparing the outcomes associated with each fMRI task, we concluded that the literature was too heterogenous; no single task was used in more than four papers, and many tasks were used in only a single paper. Even when tasks overlapped across papers, researchers did not always choose the same BOLD contrast to correlate with behavior. Assessments of risk-taking outcomes (both behavioral performance and self-report) also varied widely, with no single measure used in more than two papers. Furthermore, fewer than half of the studies reviewed utilized a whole-brain analysis; rather, most authors limited their search space through the use of either an ROI approach or a task-activation map.

Because of the highly heterogeneous nature of both fMRI tasks and individual outcome measures (as well as the preponderance of region of interest approaches), we elected not to perform a quantitative meta-analysis but rather to more qualitatively visualize the brain regions that have been repeatedly implicated in brain-behavior correlations. We describe findings discovered through whole-brain/bottom-up analyses and those identified through region of interest (ROI) analyses, as well as those identified through both approaches.

To test the evidential value of the literature (that is, to assess whether significant findings represent real effects and are not soley the result of selective reporting), we submitted findings to a p-curve analysis, a technique that tests for publication bias in a group of studies (Simonsohn, Nelson, and Simmons, 2014). Publication bias can be difficult to assess in the neuroimaging literature, because multiple non-independent results are typically reported, approaches to thresholding and correction for multiple comparisons vary widely, and often only the maximum z or p-statistics are reported in tables (David et al., 2013). However, the frequency with which ROI approaches were used in this literature allowed us to investigate publication bias as it applied to any correlations hypothesized a priori (a priori ROIs were identified in over 50% of the papers). In the present corpus, some researchers relied on correlational approaches to correlate BOLD responses with risk taking, while others utilized a regression with other covariates; a p-curve analysis allowed us to examine the results of these studies combined even though different test statistics (R, t, z, beta, etc.) were reported across studies (Simonsohn, Nelson, and Simmons, 2014; Simonsohn, Simmons, and Nelson, 2014).

Creating a p-curve involves plotting the distribution of p-values across multiple studies and investigating the shape of the resulting distribution. A p-curve for a significant effect with an unbiased distribution of p values (e.g., one that is not subject to p-hacking or other publication bias) will be right-skewed, with the majority of p-values in the lower range. A flat distribution suggests that publication bias may not be of concern, but that the effect size is small or nonexistent. A left-skewed distribution suggests that publication bias, and potentially even p-hacking, is a concern for a group of studies (Simonsohn et al., 2014). Following the technique developed by Simonsohn and colleagues, we created a p-curve disclosure table and plot for all papers that tested a brain-behavior correlation using a region of interest defined a priori. We tested the shape of the distribution using a binomial test; Simonsohn and colleagues suggest this simple method to determine whether the curve is right-skewed and therefore contains evidential value. Frequently, more than one p-value was reported (e.g. when more than one ROI or contrast was tested or when a relation was tested with and without a covariate). In these cases, we selected only the first reported p-value to maintain independence of findings. In cases where multiple p-values are reported, Simonsohn and colleagues (2013) recommend testing different versions of the analysis. Therefore, to confirm the results of our p-curve analysis, we also performed a related analysis using only the most the commonly selected ROI, the ventral striatum. This ROI appeared in nine out of twelve studies that utilized an ROI approach.

The nucleus accumbens of the ventral striatum is a sensible choice for an ROI in this literature since it has been implicated in several aspects of risk-taking behavior. Hypersensitivity of the VS during adolescence is hypothesized to contribute to the “imbalance” between affective and cognitive control systems (Casey et al., 2016; Shulman et al., 2016). A related hypothesis posits that the presence of peers heightens VS responsivity during adolescence, thereby increasing susceptibility to health risk behavior (Albert, Chein, and Steinberg, 2014; Chein et al., 2011). These theories of adolescent risk taking, which concern trajectories of normative development, are sometimes at odds with theories of aberrant brain functioning in addiction. Both theories of addiction and adolescent decision-making concern the neural substrates of reinforcement and reward, particularly the mesolimbic dopamine system. The nucleus accumbens appears to play a crucial role in the acute reinforcing effects of a variety of drugs, as well as subsequent cravings (for a review, see Koob and Volkow 2010), and some drugs, such as cocaine, have been shown to affect dopamine receptor availability (Volkow et al., 1993). However, hypo-functioning of reward circuitry has been hypothesized to predispose individuals to risk for addiction, such as in the reward deficiency hypothesis (Blum et al., 2000), although support for this hypothesis has been mixed (Hommer et al., 2011; Peters et al., 2011). Bjork and Pardini (2015) point out the comorbidity of substance use disorders and both disruptive behavior disorders and anti-social disorders, both of which have been associated with heightened reward sensitivity (e.g., Byrd et al., 2014). Thus, competing theories might suggest that adolescents who are especially susceptible to risky behavior may either show more reward sensitivity than their peers, or that such adolescents may evince relatively less reward sensitivity. Given these mulitple hypotheses, we report all significant results (positive and negative correlations) in our results. Results of the p-curve analysis for the VS/NAcc only are reported in Supplementary Materials. In addition to our discussion of whole-brain and ROI effects with respect to different measures of risk taking, we provide several more general critiques of the literature and make suggestions for best practices in future work.

## Summary of Findings

Table 1 summarizes the findings of the 23 studies we identified that correlated individual differences in neural responses with risk taking propensity among adolescents. Twenty studies investigated neurotypical youth, one investigated a population with experience with substance abuse, and two included individuals in both of these categories. While some studies also directly compared low frequency and high frequency drug or alcohol users, we focused specifically on correlational findings also reported in these papers.

### Is activation in certain brain regions related to individual differences in risk taking?

Brain regions implicated in brain-behavior correlations were highly heterogeneous and included over 25 cortical and subcortical regions (Table 1). Figure 1 depicts these brain regions and whether they were found in whole-brain/bottom up approaches, targeted ROI analyses, or both. The most commonly implicated region (combined whole-brain and ROI analyses) was the medial prefrontal cortex (broadly defined), which was related to risk taking in eleven studies, followed by VS/NAcc (nine studies), and the insula and orbitofrontal cortex (each in four studies).

**Figure 1.**
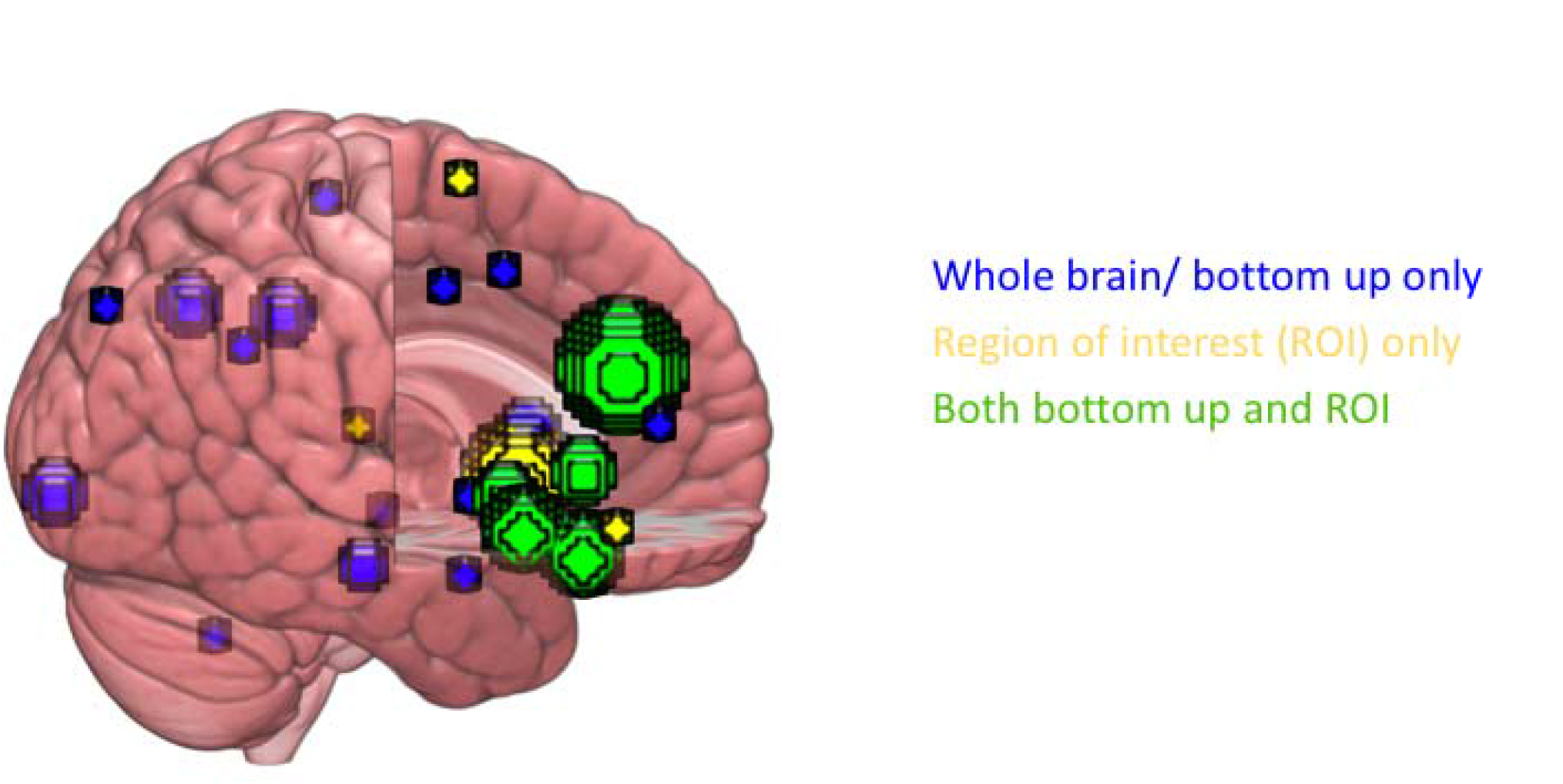
Brain regions for which significant activation during task-based fMRI was found to correlate with risk-taking behavior. Over 25 cortical and subcortical regions were implicated in brain-behavior correlations linking task-based activation to risk-taking behavior or related measures. The size of the sphere reflects the frequency with which each brain region was implicated in the literature, with larger spheres representing brain regions that were more commonly identified in brain-behavior correlations. Note that this image is a qualitative depiction of findings based on authors’ labels for brain regions (e.g., ventral striatum, mPFC) rather than a quantitative meta-analysis.

Some of the same brain regions were identified through bottom up analysis and through ROI analysis. Nonetheless, the disparity between findings that result from these two approaches is quite striking. While the VS was the most frequently hypothesized a priori ROI, activity in this region was *never* reported to correlate with risk taking in bottom-up approaches. In contrast, only two studies investigated the mPFC using an ROI approach, but activation in mPFC or vMPFC was identified as correlating with behavior in eleven out of the fourteen studies using bottom-up analyses. Several of the most commonly identified brain regions in bottom-up analyses, including the posterior cingulate cortex (PCC), the precunues, and the occipital cortex broadly defined, were never investigated a priori.

The disparity between findings discovered through bottom-up and ROI methods hints that results do not converge across studies. Rather, the overall pattern of findings suggests that that study outcomes tend to emerge specifically in the places where one goes looking. And while, at first blush, the frequency of positive findings in the VS would seem to support the notion that individual differences mirror age differences (since the VS is also often implicated as a locus of age differences), several of the most frequently implicated brain regions discovered in this review (e.g., the PCC, precuneus), are not central components of either reward circuitry or a cognitive control network, and are therefore not generally discussed in theories of adolescent decision-making. The general lack of convergence across studies may be explained by the wide variety of approaches used to elicit brain-behavior correlations, including a range of both fMRI tasks and outcomes measures used to assess risk taking, as well as the prevalence of region-specific, rather than whole-brain, analytic approaches.

### Are findings related to the particular tasks used to investigate brain-behavior correlations in adolescent risk taking?

In total, sixteen unique tasks were used to elicit brain responses during fMRI (Table 1); indeed, only five tasks were used in more than a single study: the Monetary Incentive Delay task (MID; four studies), Wheel of Fortune (two studies), the Stoplight Task/Risky Driving Task (two studies), the Balloon Analogue Risk Task (two studies), and Go/No-Go (two studies). Eleven additional tasks were used just once. Eight tasks assessed risk-taking behavior directly; all other tasks involved stimuli that were hypothesized to relate to risky outcomes in some way, such as reward processing, cognitive control, or viewing images of risky behaviors.

The most commonly utilized task was the MID, a task which has been used to characterize the neural correlates of reward response in neurotypical and clinical populations (Knutson, Westdorp, Kaiser, & Hommer, 2000). The MID contains no risk-taking behavioral component; participants win money by pressing a button after seeing a prompt. Earning money (or, on punishment trials, avoiding the loss of money) is contingent on responding within a certain window of time. However, the MID is not a decision-making task, as players only have a single option for maximizing their profits and do not have to engage in a cost/benefit analysis. Similar reward tasks were also used in other studies investigating individual differences in adolescent risk taking. Galvan and colleagues (2007) used a more youth-friendly task in which participants were asked to identify the location of a pirate cartoon by pressing a button, with correct answers earning money pots of different sizes. This task involved a choice between two options (each displayed on the left or right side of the screen), but demanded no risk assessment or gambling. Braams and colleagues (2015, 2016) utilized a reward task based on chance rather than correct responses; participants played a coin toss game for heads or tails and randomly won money based on the outcome. While the authors call the task a “gambling game,” participants do not choose whether to bet money; therefore no risky decision-making was involved. These three non-risky reward tasks were designed explicitly to elicit a response in the VS/NAcc and other reward-related regions, and all studies utilizing these tasks hypothesized a correlation between VS/NAcc response and behavior (see Table 1).

A handful of studies involved other non-risk-taking tasks designed to elicit activation in brain regions implicated in executive functioning, particularly response inhibition. For example, Cascio and colleagues (2015) utilized a Go/No-Go task and hypothesized that individual differences in risk-taking behavior (as indexed by risky driving behavior and survey measures) would be related to activation in regions implicated in inhibition, including the inferior frontal gyrus and basal ganglia. The authors used an ROI approach to investigate correlations with activation across the entire basal ganglia, a group of subcortical nuclei that include the VS/NAcc and are also implicated in reward. Mahmood and colleagues (2013) also utilized Go/No-Go, but focused on brain data extracted from the angular gyrus and ventromedial prefrontal cortex; activation in these regions on No-Go trials was related to drug use and drug and alcohol dependency symptoms, but these findings were driven by high-frequency drug and alcohol users. Chung and colleagues (2015) investigated response inhibition using an anti-saccade task, in which participants were instructed to inhibit a prepotent eye movement. The authors added a reward component to the task by incentivizing successful inhibition. Like Cascio et al., Chung and colleagues investigated correlations in *a priori* ROIs related to their task; these included brain regions involved in response inhibition and reward.

Eight fMRI tasks used in this literature did require participants to engage in some sort of risky decision-making. Six of these tasks were gambling games, in which participants either bet money or selected between options with different-sized rewards. Most gambling tasks involved binary choices (e.g., Cservenka, Jones, and Nagel, 2015; Eshel, Nelson, Blair, Pine and Ernst, 2007, Paulsen et al., 2012; Op de Macks et al., 2016, Van Leijenhorst et al., 2010), although these choices varied: participants might select outcomes with different expected values (Cservenka et al., 2015; Eshel et al., 2007; Van Leijenhorst et al, 2010), choose an uncertain or certain outcome (Op de Macks et al, 2016) or select whether or not to play a round entirely (Barkley-Levenson, et al., 2013; Paulsen et al., 2012). The task used by Barkley-Levenson and colleagues (2013) involved two gambling options (Accept or Reject a gamble), but participants could “strongly” or “weakly” select an outcome; these additional choices were used to minimize participants’ tendency to default to the same choice over and over.

Only one gambling task, “Wheel of Fortune,” was used more than once (Cservenka, et al., 2015 and Eshel et al., 2007; see also the “Cake Gambling Task,” a similar task used in Van Leijenhorst et al. 2010). In Wheel of Fortune, participants select one of two colors, each of which has a different probability of being selected and a different associated reward value. The task is divided into three phases: color selection, anticipation of outcome, and feedback. Eshel and colleagues correlated task performance with the BOLD contrast comparing trials in which participants selected the riskier choice to trials in which they selected the safe choice. Van Leijenhorst et al. (2010) focused on the BOLD contrast comparing high risk and low risk trials and added participant choice as a regressor in the model. Cservenka and colleagues used the Win > No Win contrast (during feedback) in their ROI and whole-brain investigations, and described their findings in terms of reward processing, rather than risk taking. Thus, while the paradigms used by Cservenka et al., Eshel et al., and Van Leijenhorst et al. were highly similar, they are difficult to compare directly, as each relied on different contrasts of interest for the correlational analysis. Finally, Xiao and colleagues (2012) used the Iowa Gambling Task; this task differs from those described above in the higher number of options available to the participant (they select a card from one of four decks), and in the role of feedback learning. In sum, even the gambling tasks utilized in this literature vary widely in terms of definition of risk, game structure, and contrasts used in subsequent analysis.

Two other risky decision-making tasks were used in multiple papers: the Stoplight Task/Risky Driving Task, and the Balloon Analogue Risk Task (BART). The Stoplight Task, in which participants complete a driving simulation and decide whether or not to run a series of yellow lights, has been used in several fMRI and/or behavioral studies investigating age differences in risk taking (Chein et al., 2011; Steinberg et al., 2008; Kahn et al., 2016; Voroyev et al., 2015). However, whereas Chein and colleagues (2011) used the BOLD contrast comparing brain activity at yellow lights to all other parts of the task, studies from the current literature used the contrast of Stop > Go (Kahn et al., 2016; Vorobyev et al., 2015) and/or Crash > No Crash (Kahn et al., 2015). The BART (Lejeuz et al., 2002) is a task in which participants are instructed to press a button to pump up a balloon. As the balloon inflates, participants win more money or points. However, if the balloon explodes, participants lose all of the money or points accumulated on that balloon; thus, every additional pump confers greater risk but also greater potential reward. Telzer and colleagues (2015; see also, Telzer et al. 2013a, 2013b, 2013c, Qu et al., 2015, 2016) modeled balloon pumps parametrically: they investigated which brain regions exhibited greater activation as participants continued to select the risky choice. McCormick and Telzer (2017) used the BART but instead focused on the contrast between balloon pumps following negative feedback (i.e., the balloon after a trial in which the balloon exploded) and pumps following positive feedback (i.e., the balloon after a trial in which the participant won money).

Finally, several researchers used tasks that did not involve decision-making, but rather presented complex, realistic stimuli such as photographs or videos (Pfeifer et al., 2011; Saxbe et al., 2015; Sherman et al., 2017). Pfeifer and colleagues asked participants to passively observe images of emotional faces, and Saxbe and colleagues asked participants to view silent video clips of peers and adults and imagine what the individuals were feeling at the moment. Sherman and colleagues (2017) presented photographs of risk-taking behavior or paraphernalia (e.g. teens drinking alcohol or a marijuana pipe) in a task mimicking the social media app Instagram, and asked participants to decide whether or not to “Like” each image.

The reliance on analytic approaches with restricted search spaces (e.g. ROI analyses or searches within task-activation maps) precluded us from performing a quantitative analysis investigating patterns of brain-behavior correlations as a function of task choice. Indeed, areas in which correlations with behavior were found tended to be areas that also showed a main effect of task. For example, Saxbe and colleagues (2015) and Sherman and colleagues (2017) both presented participants with realistic social stimuli and found that self-reported risk taking was correlated with response in the precuneus and the PCC, brain regions showing main effects of task and also implicated more generally in social cognition. Similarly, all of the studies using the MID, a task designed to elicit a reward response in the brain, reported a correlation with behavior in at least one part of the striatum. Because papers described here limited their search to either *a priori* ROIs known to be active during the fMRI task (e.g., Braams et al., 2016, Bjork et al., 2008), or performed bottom-up analysis constrained within a task activation map (e.g. Benningfield et al., 2014; Sherman et al., 2017), it appears likely that the heterogeneity of brain regions implicated in this literature is at least in part the result of both the fMRI paradigm and the use of targeted analysis.

### How is the outcome measure of “risk taking” defined?

Even greater than the number of fMRI tasks used to elicit a brain response was the number of measures used to index risk-taking behavior or propensity: over 30 different measures were utilized (Table 1). Many studies used more than one measure of risk taking. Broadly, the outcome measures can be categorized as either lab-based or real-world. Lab-based measures included any sort of risk-taking behavior assessed during the task itself, such as behavior on the Stoplight Task, delay discounting, various gambling tasks, and the BART. Real-world measures included surveys designed to assess one or more of the following: actual experiences with drug and alcohol use (e.g. CARE-R; Katz, Fromme, and D’Amico, 2000), problem behaviors associated with drug and alcohol use, (e.g. the AUDIT, Babor, Higgins-Biddle, Saunders, Monteiro, 2001) perception of the potential risks or benefits associated with activities (e.g. Benthin Risk-Perception Scale, Benthin et al., 1993), and experiences with non-drug related risk behavior and general sensation-seeking (e.g. the BSSS, Hoyle et al., 2002).

While the measures used vary widely, it is also likely that several would be correlated within a sample. For example, Sherman and colleagues (2016) reported that the sections of the CARE-R measuring perception of risk were correlated with the section assessing frequency of risk-taking behavior. Meta-analyses have also revealed correlations between sensation-seeking and alcohol use (Hittner and Swickert, 2006) and between alcohol use and risky sexual behavior under certain circumstances (Leigh 2001; Tapert et al., 2001). Nonetheless, it is also most certainly the case that the behaviors indexed by these measures differ somewhat in their neural correlates. The *opportunity* that individual adolescents are afforded to engage in drug and alcohol use or sex undoubtedly plays a role in the extent to which self-report measures like sensation-seeking and impulsivity predict risky behavior (Shulman et al., 2016).

### Are lab-based and real-world measures of risk taking correlated?

Within the current literature, we examined whether reports of real-world risk-taking behaviors, as assessed by survey measures, were related to performance on lab-based measures. Significant correlations would be in line with past research in non-fMRI settings that has documented the validity of tasks such as the IGT (Dahne et al., 2013), Stoplight Task (Centifani et al., 2014, Kim-Spoon et al., 2015), and the BART (Aklin, Lejuez, Zvolensky, Kahler, and Gwadz, 2005; Fernie, Cole, Goudie, and Field, 2010; Hansen, Thayer, and Tapert, 2014). Eleven out of the twenty-three studies on our list reported both lab-based risk taking and “real-world” measures of risk-taking behavior - that is, survey measures of attitudes toward or frequency of risky behavior, or (when groups were also defined) categorical comparisons of risky and non-risky youth. Surprisingly, however, these measures were rarely correlated. Of the eleven studies reporting both kinds of measures, only two reported a significant relation between real-world and lab-based risk taking, and two reported limited or equivocal convergence between measures (see Supplementary Table 1 for more details). Seven studies failed to find any evidence of a correlation. These findings were surprising given the successful efforts described above to establish the external validity of the lab measures. It is possible that many of the samples in the current literature were underpowered to detect such correlations, a concern that is also at least as relevant, if not more so, for the detection of brain-behavior correlations.

### Does the literature show evidence of publication bias?

As considered above, the heterogeneity of findings may be due primarily to the variety of tasks, measures, and ROIs used to investigate brain-behavior correlations; each task and measure is certain to capture a different aspect of “risk taking.” It also possible that the presence of Type I and Type II errors have contributed to mixed findings. Small sample sizes in fMRI studies can contribute to Type II errors (Dubois and Adolphs, 2016; Button et al., 2013; Yarkoni & Braver, 2012; Yarkoni, 2009). Yarkoni (2009) suggested that, in a correlational fMRI study, a sample of 50 participants has only a 66% chance of detecting a correlation of .5 at p < .001. Dubois and Adolphs (2016) recommend a sample size of greater than 100 for correlational studies. As reported in Table 1, the sample sizes in the majority of the studies we reviewed are smaller than these recommended numbers, and often substantially so. Only seven studies included 50 or more adolescent participants, and only two of these included a sample size greater than 100. Based on the sample sizes in the current literature, it is reasonable to assume that the prevalence of Type II errors is of concern, and may explain in part the limited overlap in findings. We would note, however, that many of the studies in the current corpus reported brain-behavior correlations as a secondary analysis following an exploration of main effects; thus, it is likely that sample sizes were determined based on power calculations for main effects rather than correlations.

Might false positives be an issue in the literature as well? In order to investigate a potential source for systematic false positives, we conducted two p-curve analyses, the first on the first reported correlational ROI finding in each paper utilizing ROI analyses, and the second on the most commonly reported ROI, the ventral striatum. As both analyses contain overlapping information, findings for the VS p-curve are reported in Supplementary Materials; however, the conclusions are the same. Thirteen papers tested the hypothesis that activation in least one ROI was related to risk-taking behavior, either in the lab or as reported on survey measures. Details of the statistics used to develop the p-curve are presented in Table 2, and the p-curve chart is presented in Figure 2. Simonsohn and colleagues (2013) suggest that in order for a result to show evidential value - that is, in order for us to rule out the possibility that selective reporting is solely responsible for the significant effects observed in the literature - the p-curve should be right-skewed. Furthermore, in a literature with evidential value, a binomial test should reveal that significantly more than half of the p values are < .025. Our results do not support the conclusion of evidential value. While a slight right skew appears to be evident in Figure 2, seven out of thirteen p values are > .025, and the results of both binomial tests were nonsignificant (*p* = .71 for the first reported ROI and *p* =.96 for the VS/NAcc). These results suggest that correlations (both generally and in the VS) may have been subject to selective reporting, or that the actual effect size for the correlational phenomenon under investigation is negligible. The fact that neither p-curve was left-skewed, however, favored the interpretation that effect sizes for region-of-interest findings are negligible (Simonsohn et al., 2014).

**Figure 2.**
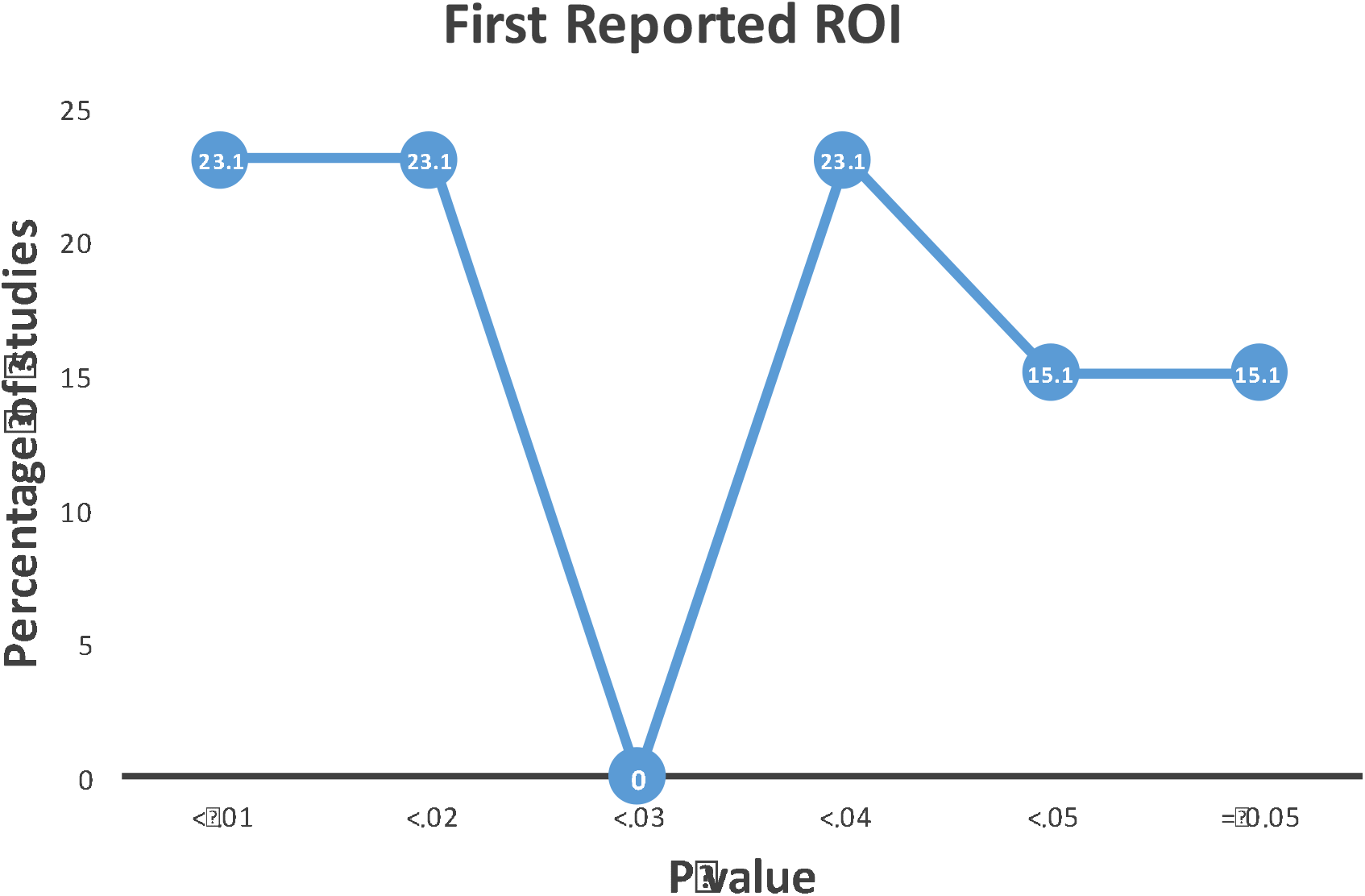
Results of a p-curve analysis for the first reported region of interest (ROI). Thirteen studies reported significant correlations between risk-taking behavior and activation in at least one hypothesized ROI.

**Table 2.**
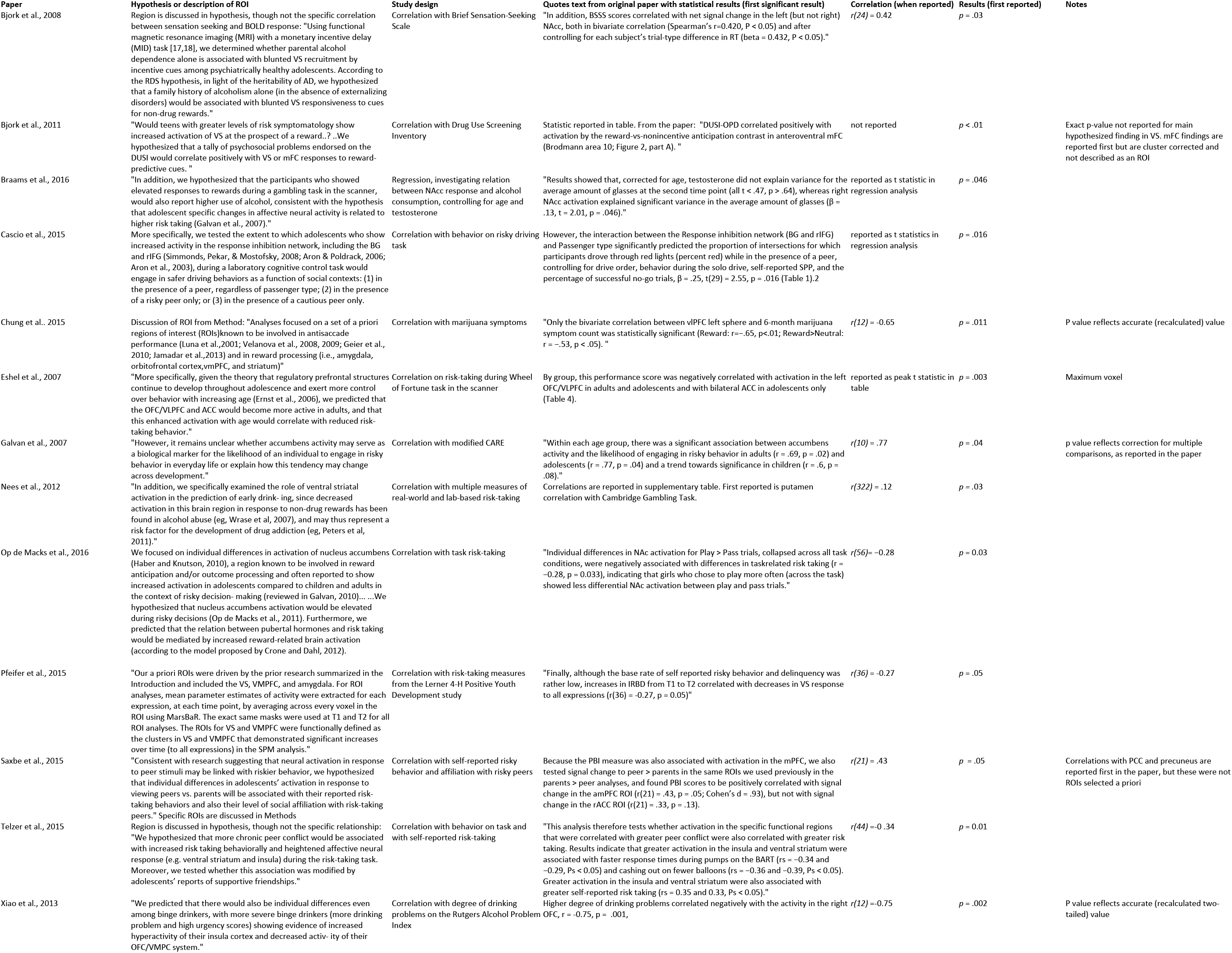
Results of p-curve analysis for first reported region of interest (ROI) analysis

While the results of our p-curve analysis leave open the possibility of publication bias, we do not find indicators of the presence of any deliberate attempts to misrepresent findings on the part of the authors. Indeed, the opposite is true, as many authors reported nonsignificant as well as significant findings. For example, Braams and colleagues (2016), in their longitudinal study relating reward response to frequency of alcohol use, reported that NAcc activation explained significant variance in the average amount of glasses of alcohol drunk per night at Time 2, but also reported several nonsignificant relations. These non-significant findings, which are not considered in the p-curve analysis, provide yet further support for the conclusion that effect sizes under consideration are small or non-existent. P-curve analyses can only be conducted on significant findings (Simonsohn et al., 2014), so the nonsignificant findings are not represented in the test of bias. Although authors may do their best to report all findings, significant and otherwise, readers may recall only significant findings in a study, and these findings can be cherry-picked to support subsequent theories or conclusions. Unfortunately, the results of our p-curve analysis suggest that there is little if any evidential value to the hypothesis that activation in individual brain regions, particularly the VS/NAcc, is correlated with risk-taking behavior. That is, it is not possible to rule out that these significant findings, of which there are many, represent anything more than selective reporting or spuriously discovered correlations. Future correlation work with sufficiently-powered samples may provide evidence to suggest otherwise, but at this time, it would be inappropriate to conclude that the literature has revealed robust one-to-one correlations between risk taking and activation in a brain region of interest.

## Discussion, Future Directions, and Recommendations

In the years since Pfeifer and Allen (2013) and Bjork and Pardini (2015) called for a greater focus in adolescent neuroscience on individual differences in risk taking, the size of the literature has increased substantially. We were surprised to discover, however, that no brain region has been consistently correlated with risk-taking behavior; nor have the implicated regions comprised a single network or system. No single brain region was implicated in more than 50% of the studies, and a p-curve analysis revealed no evidential value for the most commonly reported ROI, the VS, or for ROI findings as whole. Furthermore, the VS was never identified as correlated with behavior in whole-brain analyses.

On the one hand, these findings may seem disappointing: with such limited convergence, how are we to make informed decisions about which methodological approaches are most effective or appropriate? On the other hand, we suggest that this limited convergence actually may indicate that neural predictors of adolescent risk taking are not localized, but, rather, that explanatory power may be drawn from studies of activation across the brain. Efforts to home in on individual brain regions or even networks, while understandably motivated by theories established in the developmental literature, may nonetheless be discarding valuable information available about the rest of the brain. Furthermore, we posit that the wide range of fMRI tasks and measures of risk behavior or propensity highlight a need for further refinement of individual research questions. Thus, we have several suggestions for the field moving forward, as well as open questions that we hope can be addressed in both ongoing discussion and empirical inquiry.

Best practices for identifying the neural underpinnings of risk-taking behavior will not be one-size-fits-all, but will rather depend on the precise definition of each of these concerns.. We therefore propose that researchers wishing to investigate brain-behavior relations on the topic of adolescent risk taking should precisely define four aspects of the research question and goal: (1) the relevant population(s), (2) the hypothesized brain system or systems, (3) the hypothesized psychological factor(s) relating brain responses to outcomes, and (4) desirable and undesirable outcomes. Below, we make specific recommendations based on the definitions in these four categories.

### Defining the population

First, we must consider which population is most relevant: do we expect that neural “risk factors” exist on a spectrum ranging from typically developing individuals to those with substance use disorders or other psychiatric disorders (e.g. conduct disorders)? This question has implications for the population(s) recruited for a study, and for the decision to examine outcomes categorically or continuously. Certainly, neural systems that have been proposed to account for heightened risk taking in adolescence are also those hypothesized to confer risk for drug and alcohol addiction, including circuitry implicated in reward and reinforcement, and in executive function. It has not been established, however, whether individuals prone to either addiction or dangerous levels of sensation-seeking are simply on the further end of a spectrum that is linked to adolescent risk taking in general. Correlational approaches like those considered in the present review are appropriate if one expects the nature of neural mechanisms to be continuous. However, if “high risk-takers” are instead a qualitatively different group (e.g. as proposed by Bjork and Pardini, 2015), a categorical approach would be more appropriate. Categorical comparisons have made up the majority of studies in the clinical literature (Heitzeg et al., 2015; Squeglia and Cservenka, 2017).

In defining a sample and considering recruitment strategies, we recommend that researchers also attempt to differentiate between *effects* that risks like drug and alcohol use have on the brain and neural factors that *predispose* some individuals to make risky choices. In order to do this, a longitudinal approach that uses brain responses at one point in time to predict future outcomes is necessary. These prospective designs are increasingly common in the clinical literature (Dager et al., 2014; Heitzeg et al., 2014; Norman et al., 2011; Squeglia et al., 2016; Wetherhill et al., 2013), and we are encouraged by recent efforts to take this approach in the presently reviewed literature (e.g. Braams et al. 2016, Chung et al., 2015; Cservenka et al., 2015), as well as in recent NIH-funded efforts to investigate vulnerability in longitudinal samples.

### Defining hypothesized brain mechanisms

If we, as developmental cognitive neuroscientists, want to cite “preventing adolescent risk taking” as a motivation for grant proposals that focus on neurotypical youth, we should also be able to articulate a compelling case for how studying normative brain development will help us to address this issue. To do so, we must be specific about the neural mechanisms proposed to underlie risk taking. Researchers should clearly describe which brain systems or processes are expected to be related to risk-taking behavior, and in what way (e.g. through activity in a single region or connectivity between regions).

### Regional approaches to defining hypothesized brain mechanisms

Much of the extant literature has been successful in defining specific hypothesized brain mechanisms, likely as a result of the popularity of intuitive brain-based theories of adolescent decision-making and addiction.The frequency of ROI analyses in this literature, for example, highlights the hypothesis-driven nature of inquiry in the field, and also effectively sidesteps criticisms about reverse inference as well as ongoing concerns about cluster correction for multiple comparisons (Eklund et al., 2016). Nonetheless, our findings suggest that brain regions frequently implicated in mentalizing (mPFC, precuneus, PCC) may deserve greater targeted attention based on their frequent appearance in the whole-brain literature. Furthermore, researchers must be cautious about using ROI approaches only, as our p-curve analysis suggests that these approaches may lack evidential support. We recommend that, when researchers hypothesize that activity in a particular brain region has predictive power, they perform an ROI analysis, but also make whole-brain results available -- ideally in the form of full, unthresholded brain maps uploaded to a public repository -- for subsequent meta-analyses.

### Circuit-based approaches to defining hypothesized brain mechanisms

We must think carefully not only about where we measure responses in the brain, but the *way* in which we do so. The majority of the literature has investigated level of “activation” by extracting parameters from individual BOLD contrasts generated through classic parametric approaches. This strategy is sensible if one expects that the magnitude of response in specific regions will provide the most explanatory power, perhaps because this brain region is especially “sensitive” or “efficient” in some individuals (though see Poldrack, 2015). However, current theories of adolescent risk taking do not necessarily assume that this is the case. For example, Casey, Getz, and Somerville (2016) recently called for a shift away from region-based or node-based approaches to the study of brain development and instead encourage a consideration of brain development with respect to brain circuitry, particularly when it comes to the interaction of regions involved in affect and motivation with those involved in cognitive control. Accordingly, a functional or dynamic connectivity approach might be a more appropriate way to explain individual differences. Some examples can be found in this literature already, including studies exploring psychophysiological interactions (Bjork, Knutson and Hommer, 2011, Pfeifer et al., 2011; Qu et al., 2015). Of course, the recommendation to consider connectivity approaches does not suggest that these approaches are free from limitations, only that the analytic approach should reflect the hypothesized nature of the brain mechanisms.

### Multivariate, predictive approaches to defining hypothesized brain mechanisms

An alternate approach that could prove fruitful in this domain might involve abandoning the notion that the properties of individual brain regions, or even brain networks, can provide a window into individual differences in risk behavior, and to turn instead turn to analytic techniques in which parameters from the entire brain are used to predict outcomes. Here, we can utilize approaches that are increasingly common in the clinical literature. For example, Squeglia and colleagues (2016) utilized a random forest classification model approach that predicted alcohol initiation status by age 18 in a longitudinal sample of adolescents with 74% accuracy. The researchers included a wide range of predictors, including neurocognitive and neuropsychiatric assessments, sociodemographic data, and both structural and functional parameters extracted from across the entire cortex. They found that the MRI variables contributed additional explained variance over and above the other variables. Importantly, this analytic approach modeled both main effects and interactions between variables, including aspects of brain structure and function; these interactions also contributed to better model fit.

Predictive model approaches have the potential to provide a direct connection between research and application, especially when combined with longitudinal designs, as in the case of Squeglia et al. (2016; see also Whelan et al., 2015). For researchers specifically interested in predicting health outcomes, this approach is sensible. However, cognitive neuroscientists typically are not only interested in preventing problematic outcomes, but also in understanding the developing neural architecture of decision-making. Models that derive predictions from many brain regions make interpretation of cognitive mechanisms difficult, especially given that BOLD activation reflects not only differences in brain activity but also differences in vasculature (Poldrack, 2011). The appeal of imbalance and dual systems models is that they allow us to bridge psychological science and neuroscience by connecting cognitive processes to brain responses.

One further way that researchers might bridge psychological and neural science using whole-brain models would be to test the predictive power of different fMRI tasks in the same population. For example, does a risk taking task allow for more accurate prediction than a task that only tests reward responses or response inhibition? Squeglia and colleagues (2016), for example, were able to successfully predict initation of heavy alcohol use in a longitudinal sample using a working memory task. We encourage researchers to take a similar approach using reward and risk taking tasks. The Wheel of Fortune task may be an especially good candidate for this approach, as it includes separable selection and feedback phases, each of which could be entered into a predictive model and compared with respect to their predictive power (or combined, potentially providing additional explanatory power). The fact that researchers often have differed on the contrasts of interest selected within the same task suggests that separable phases of task performance may provide unique insight into individual differences in decision-making. Furthermore, the Wheel of Fortune is one of only a handful of tasks that have been used multiple times in this literature.

Support for the utility of whole-brain multivariate analysis combined with fMRI task comparison comes from a recent paper by Rudolph and colleagues (2017), who used whole-brain connectivity during three phases of a Go/No-Go task to investigate risk predilection (as assessed by the Benthin Risk Assessment). The authors utilized functional connectivity during a neutral Go/No-Go task to successfully predict age in a novel test sample. They then compared functional connectivity during *emotional* Go/No-Go tasks (involving the threat of an aversive stimulus or the anticipation of a reward) to neutral Go/No-Go. Individuals whose predicted age during the emotional contexts was younger than their age during the neutral context provided riskier responses on the Benthin than the emotionally “older” participants. Although this work did not use whole-brain responses to *predict* risk outcomes, it nonetheless presents a potential avenue for identifying individuals with heightened vulnerability. The use of fMRI tasks, rather than resting state data, allows researchers to interpret findings within a cognitive theoretical framework even when making predictions using data from the entire brain.

Of course, our consideration of the potential value of multivariate predictive approaches is not intended as a repudiation of current correlational approaches. Indeed, given that the sample sizes in the available literature are often insufficient to detect real correlational relationships, it may also be the case that we simply do not yet have a body of literature capable of producing far-reaching conclusions. These suggestions, therefore, can be considered as a way to extend current methods, rather than an as a replacement of region-of-interest or bottom-up approaches that focus on individual brain regions.

### Defining hypothesized psychological mechanisms

When taking a cognitive neuroscience approach to understanding adolescent risk taking, one should propose specific hypotheses not only about the neural architecture, but also the psychological mechanisms linking brain responses to real-world outcomes. For example, if we expect reward functioning to predispose some adolescents to engage in more risk taking, what do we believe explains this link? Do some teens have heightened sensitivity to peers? Do certain aspects of personality (e.g. sensation-seeking) have specific neural correlates in the ventral striatum or other brain regions implicated in reward? Does neurochemical functioning of this circuitry affect the way some individuals respond to drugs and alcohol? Each of these hypotheses would suggest the inclusion of additional measures (e.g., the Resistance to Peer Influence Scale, the Sensation-Seeking Scale, and the Alcohol Use Disorders Identification Test for the three questions above). We recommend that if researchers hypothesize that deleterious outcomes such as drug abuse and accidents are indirectly related to neural functioning, that they include a both a measure of psychological phenomenon hypothesized to connect the two as well as a measure of actual risk taking in the participants’ daily lives. Several researchers utilized both types of measures (e.g. Kahn et al., 2016; Galvan et al., 2007; Xiao et al., 2013), but rarely are psychological outcomes modeled as mediators between brain response and real-world behavior (alternatively, the brain response could be hypothesized to link the psychological phenomenon to experiences in the real-world). A notable exception is a recent paper by McCormick and Telzer (2017), which found that behavioral sensitivity to feedback (as indexed by responses on the BART) was indirectly related to real-world risk taking (as indexed by the Adolescent Risk taking Scale) though mPFC response to feedback.

In the present review, we found that laboratory tasks designed to measure psychological phenomena rarely correlated with survey and psychometric measures of purportedly similar constructs. This finding is somewhat at odds with the broader literature that successfully links performance on tasks like the BART and IGT to real-world outcomes. It is possible that low power contributes to the lack of significant findings. Nonetheless, we suggest that, when possible, researchers include both a behavioral measure and psychometric measure of the relevant psychological phenomenon if both are of interest, rather than making assumptions that one is correlated with the other just because this relationship has been reported in past work. Several of the psychometric measures we have described (BSS, RPI) consist of fewer than fifteen questions and would not add significantly to study length, especially if the behavioral task data is collected as a part of the fMRI scan.

### Defining desirable and undesirable outcomes

If the neuroscience of adolescent risk taking is to converge on a point of applicability, we (as a field and as individual researchers) need to more precisely define desirable and undesirable outcomes for youth, and to consider how these definitions are implied in our choice of measures. This issue is of particular concern when researchers correlate magnitude of response in a brain region with a measure of risk-taking behavior. Some measures (e.g. the Adolescent Risk taking Scale, CARE) simply “add up” the frequency of different risk-taking behaviors, whether for one kind of drug, all drug and alcohol use, or across many domains (e.g. playing contact sports, sexual risk taking, etc.). If complete abstinence from risky behavior is considered an ideal outcome, then this approach may be sensible. However, some amount of risk taking is normative and likely adaptive in adolescence, even when it comes to experimentation with drugs and alcohol (Shedler & Block, 1990). Researchers may therefore wish to use measures that index not only risk-taking behavior but also problematic outcomes associated with these behaviors, such as addiction symptomology, poor health outcomes, and interference with daily life (e.g. the AUDIT, Rutgers Alcohol Problem Index, Customary Drinking and Drug Use Record).

Beyond the hypothesized “desirable” or “undesirable” outcome, the choice of a risk taking measure may raise statistical concerns as well. Lebreton and Palminteri (2017) describe the ways in which floor performance on a behavioral measure can in effect reverse the direction of brain-behavior correlations, due to low-performing individuals having a smaller standard deviation (SD) than high-performing individuals on the task-related variable. This difference in SD results in statistical dependency between trait performance and the SD of the behavioral measure. Concerns about floor effects are relevant for samples with low levels of risky behavior (as is often the case with neurotypical adolescents who are able and willing to participate in fMRI research). If few participants in a sample engage in substance use, for instance, a measure that assesses only substance use (as opposed to a wider variety of risk behaviors) will capture only limited variability in the sample. Alternatively, researchers can select a subsample of participants on the basis of one or more relevant characteristics; for example, Goldenberg and colleagues (2013) specifically investigated brain and behavioral correlates of sexual risk taking, and therefore included only participants who reported being sexually active (participants were a subsample of those from Telzer et al., 2015, reviewed in Table 1).

### General suggestions for best practices

In the past year, several papers have been published that discuss the efficacy of current approaches to the study of individual differences in behavior and the brain; these papers make best practice recommendations that are applicable to cognitive neuroscientists generally (e.g., Calhoun et al., 2017; Dubois and Adolphs, 2016; Lebreton and Palminteri, pre-print). We will not consider these general recommendations at length but do remind our readers of two concerns that can completely undermine an otherwise well-designed study. First, sample sizes of fewer than 50 (and likely even 100) participants may simply be too small to reliably detect correlations between variables. While the recommendation to increase sample sizes is simple in theory, it is much more difficult in practice, given the difficulty of recruiting developmental samples and the tremendous expense of fMRI work. We are hopeful that recent efforts by the NIH to fund large multi-site studies in developmental populations (e.g. the Adolescent Brain and Cognitive Development Study and The Developing Human Connectome Project) will mitigate this issue in years to come. Similarly, longitudinal work is needed to differentiate between the effects of risky behavior (especially drug and alcohol abuse) on brain development and neural factors that predispose individuals to engage in risky activities. We are encouraged that several of the more recent papers discussed in this review utilized longitudinal designs and are hopeful that we are witnessing the beginning of a trend, especially given that the aforementioned NIH projects employ longitudinal designs.

Finally, we encourage researchers to “think outside the brain” and consider the interaction of effects at the neural and environmental level, along with genetic information. It is highly unlikely that our coarse measures of brain activity can explain the majority of the variance in risk-taking behavior, or that a single neural phenotype will always predict greater risky behavior. We therefore advocate for the use of more complex models that account for brain and environmental factors (e.g. Squeglia et al., 2016; Whelan et al., 2014). Alternatively, researchers may consider conducting case studies of small populations (e.g. a school or peer group) to document brain-behavior links in youth within highly homogenous environments. This latter recommendation may be especially suitable as an avenue for the development of targeted preventative programs.

As we move forward, we must also be mindful of ethical considerations inherent in the identification of individuals at heightened risk for suboptimal outcomes. In particular, while prospective identification of vulnerable individuals could lead to the development of targeted preventative measures, it could also open the door for stigmatization, discrimination, or a false sense of security or worry. Thus, we encourage members of the field to carefully consider not only best methodological practices, but also to begin a dialogue regarding best practices for conscientious and responsible application of research findings.

### Conclusions

The past ten years of developmental cognitive neuroscience research have produced an expansive literature on adolescent decision-making. Over these ten years, and particularly in the past three, researchers have attempted to leverage our knowledge of adolescent brain development to understand brain mechanisms that predispose some youth to engage in more risky behavior than their peers. While this line of inquiry could potentially lead to targeted preventative measures and the subsequent reduction of addiction, injury, and death in this population, there is little evidence to suggest that correlational approaches have yielded reliable, applicable findings. Though the literature we have reviewed does not provide converging evidence that singles out specific brain regions or systems as especially promising targets for research on risk -aking, we nonetheless hope that this review will serve as a snapshot of the current state of the field that will help to shape future investigations. Furthermore, we believe that progress can be made with the development of more precise research questions that move beyond a general interest in adolescent risk taking, as well as methodological approaches that move beyond one-to-one correlations between behavioral indices and activity within individual brain regions.

